## supplement for "Targeting PRMT3 Impairs Methylation and Oligomerization of HSP60 to Boost Anti-Tumor Immunity by Activating cGAS/STING Signaling"

**Supplementary Figures**

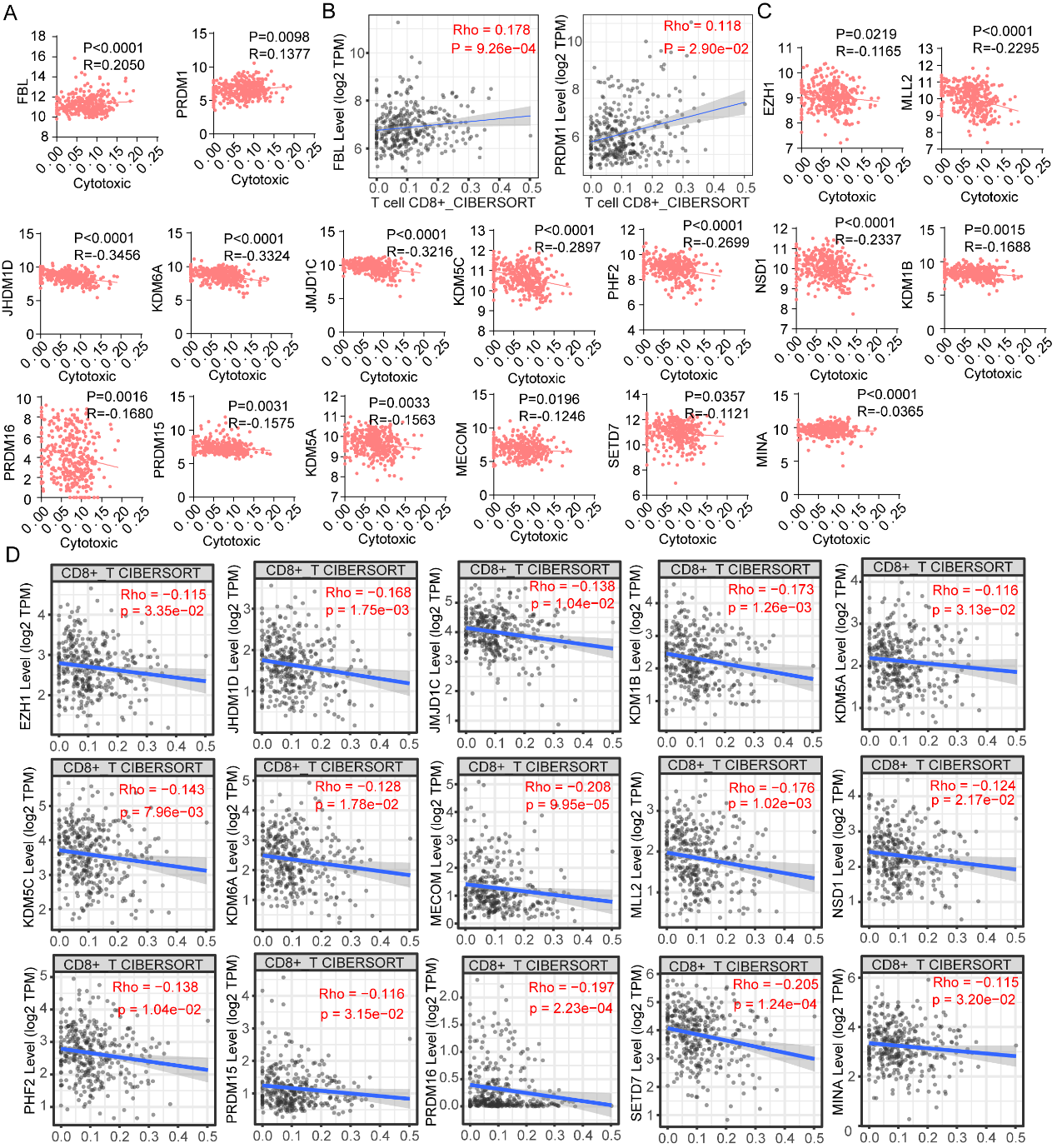

**Supplementary Figure 1**. High PRMT3 expression was strongly associated with poorer response to immunotherapy in HCC patients

A. Positive correlation between PTM expression and cytotoxic T cells score evaluated by ImmuneCell AI in TCGA-LIHC dataset.

B. Positive correlation between PTM expression and CD8+ T cell infiltration evaluated by CIBERSORT in TCGA-LIHC dataset.

C. Negative correlation between PTM expression and cytotoxic T cells score evaluated by ImmuneCell AI in TCGA-LIHC dataset.

D. Negative correlation between PTM expression and CD8+ T cell infiltration evaluated by CIBERSORT in TCGA-LIHC dataset.

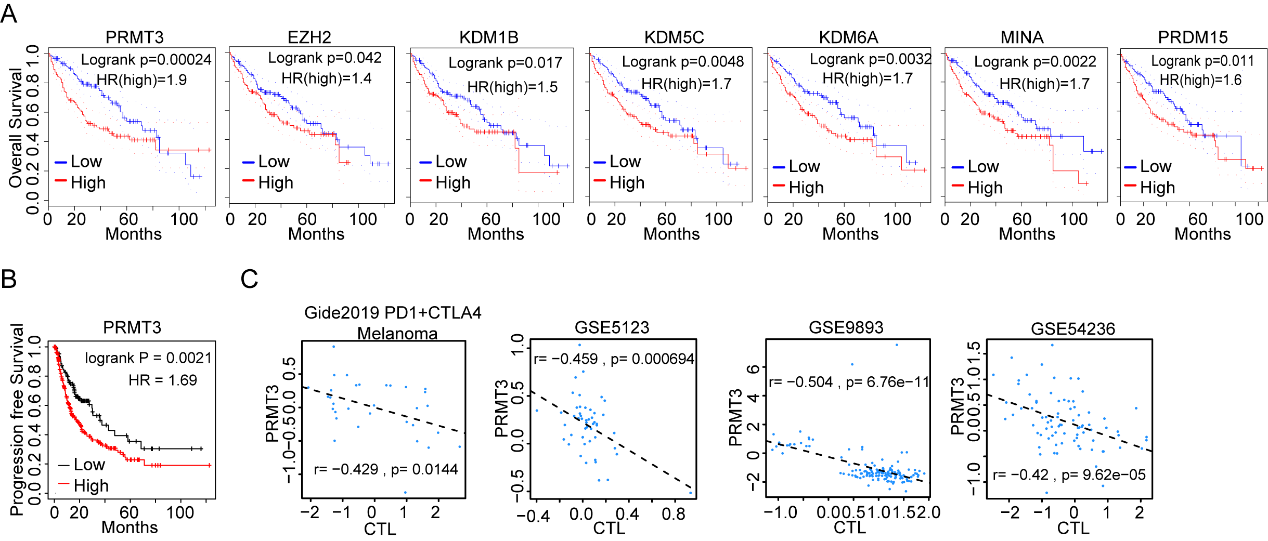

**Supplementary Figure 2**. High PRMT3 expression was strongly associated with poorer response to immunotherapy in HCC patients

A. Kaplan–Meier overall survival curves of individuals with different PTM expression in TCGA-LIHC dataset.

B. Kaplan–Meier progression-free survival curves of individuals with different PRMT3 expression in TCGA-LIHC dataset.

C. PRMT3 was negatively correlated with immune infiltration in several public dataset.

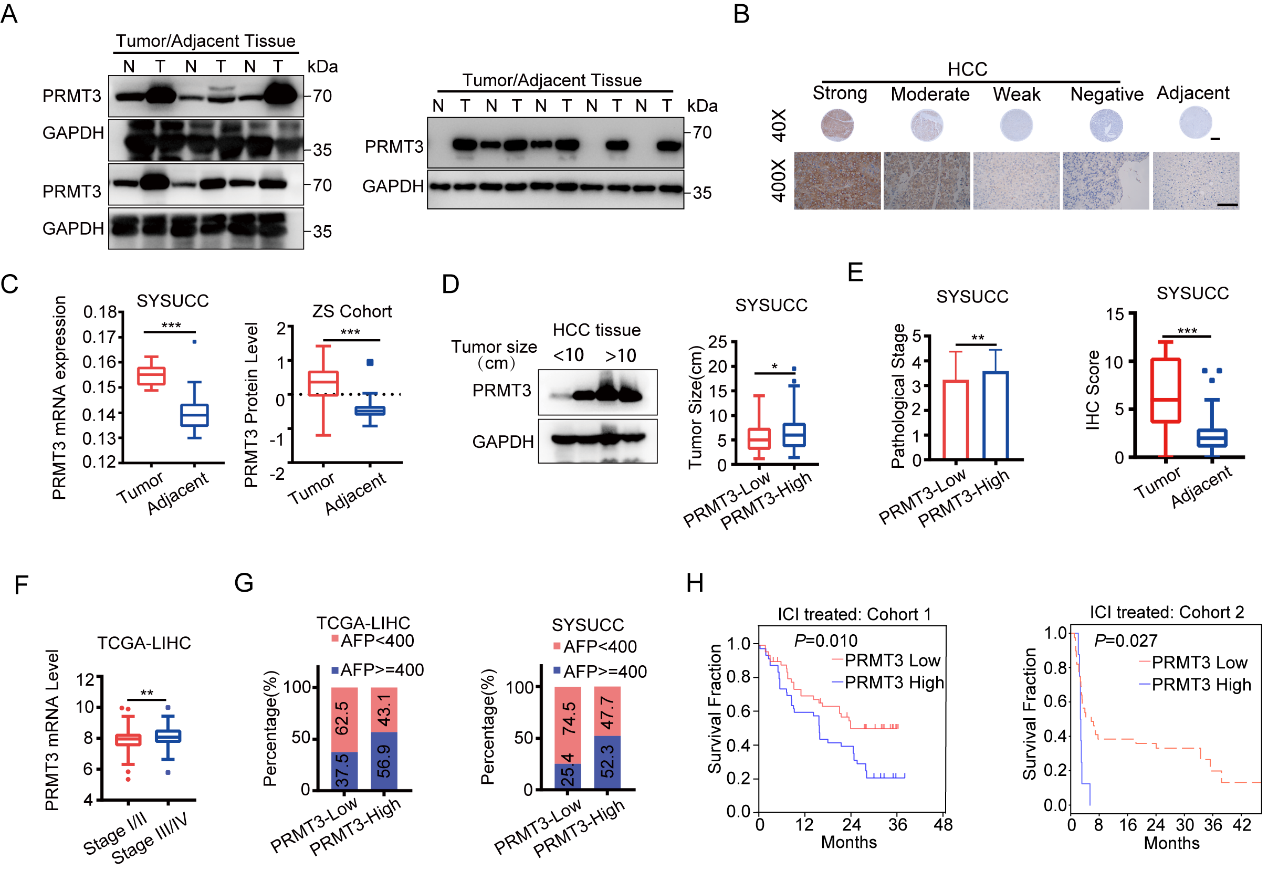

**Supplementary Figure 3. PRMT3 was up-regulated in HCC and predicts a poor prognosis**

A. PRMT3 expression was detected by western blot in HCC and adjacent tissue.

B. PRMT3 expression was detected by IHC in HCC and adjacent tissue.

C. PRMT3 expression was up-regulated in the SYSUCC cohort and a public dataset.

D. PRMT3 expression was detected by western blot and IHC in different sizes of tumors.

E. PRMT3 expression was detected by IHC in HCC of different pathological stages.

F. PRMT3 expression was detected by IHC in HCC of different TNM stages.

G. The correlation between PRMT3 and AFP expression levels.

H. Associations of PRMT3 expression with overall survival in public immunotherapy datasets.

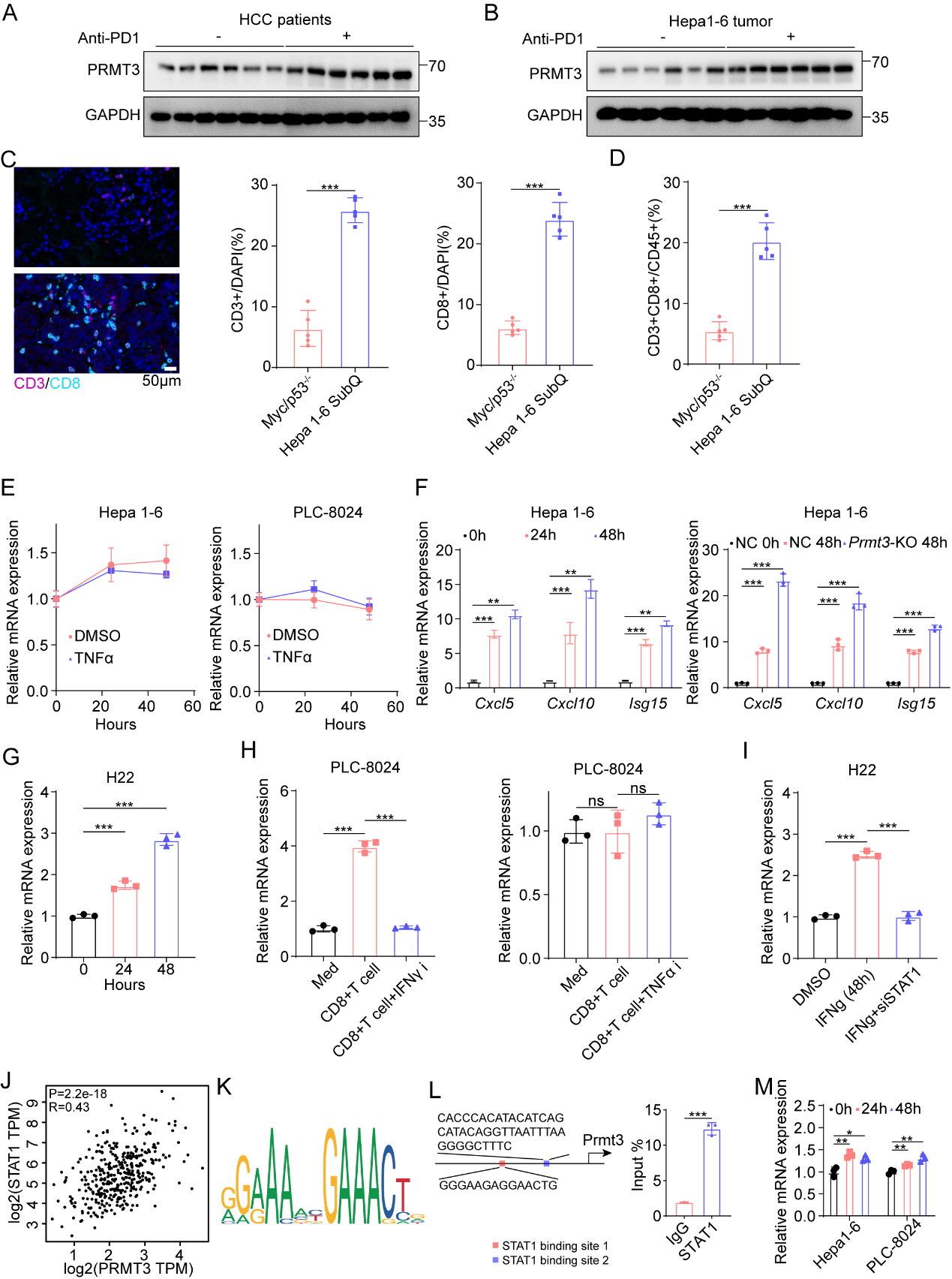

**Supplementary Figure** **4.** **Effector T cells up-regulate PRMT3 expression via IFN-γ-STAT1-dependent pathways**

A. PRMT3 expression was detected by western blot in HCC patients who received anti-PD1 therapy or not.

B. PRMT3 expression was detected by western blot and qPCR in Hepa1-6 subcutaneous tumors which received anti-PD1 therapy or not.

C. Immunofluorescence identifying CD3+ and CD8+ immune cells in subcutaneous tumors and Myc/TP53^-/-^ spontaneous tumors. Scale bar, 50 μm.

D. Flow cytometry analysis assessing the percentage of CD8+ T cell from tumors in indicated groups. Data was analyzed by FlowJo.

E. Hepa1-6 and PLC-8024 cells were treated with TNFα. PRMT3 expression was analyzed by qPCR.

F. Several interferon-stimulated genes were detected by qPCR in Hepa1-6 cells treated with IFNγ.

G. H22 cells were treated with IFNγ. PRMT3 expression was analyzed by qPCR.

H. PLC-8024 cells were treated with the conditioned medium of CD8^+^ T cells sorted from HCC tumors and inhibitors of IFNγ and TNFα. PRMT3 expression was analyzed by qPCR.

I. PRMT3 expression in *Stat1*-KD and control H22 cells as shown by qRT-PCR.

J. Correlation analysis between STAT1 and PRMT3 expression in TCGA-LIHC dataset.

K. The binding motif of STAT1 predicted by JASPAR.

L. ChIP-qPCR was used to determine the binding of STAT1 to *Prmt3* promoter region in Hepa1-6 cells treated with IFNγ.

M. Several interferon-stimulated genes were detected by qPCR in Hepa1-6 cells treated with IFNα.

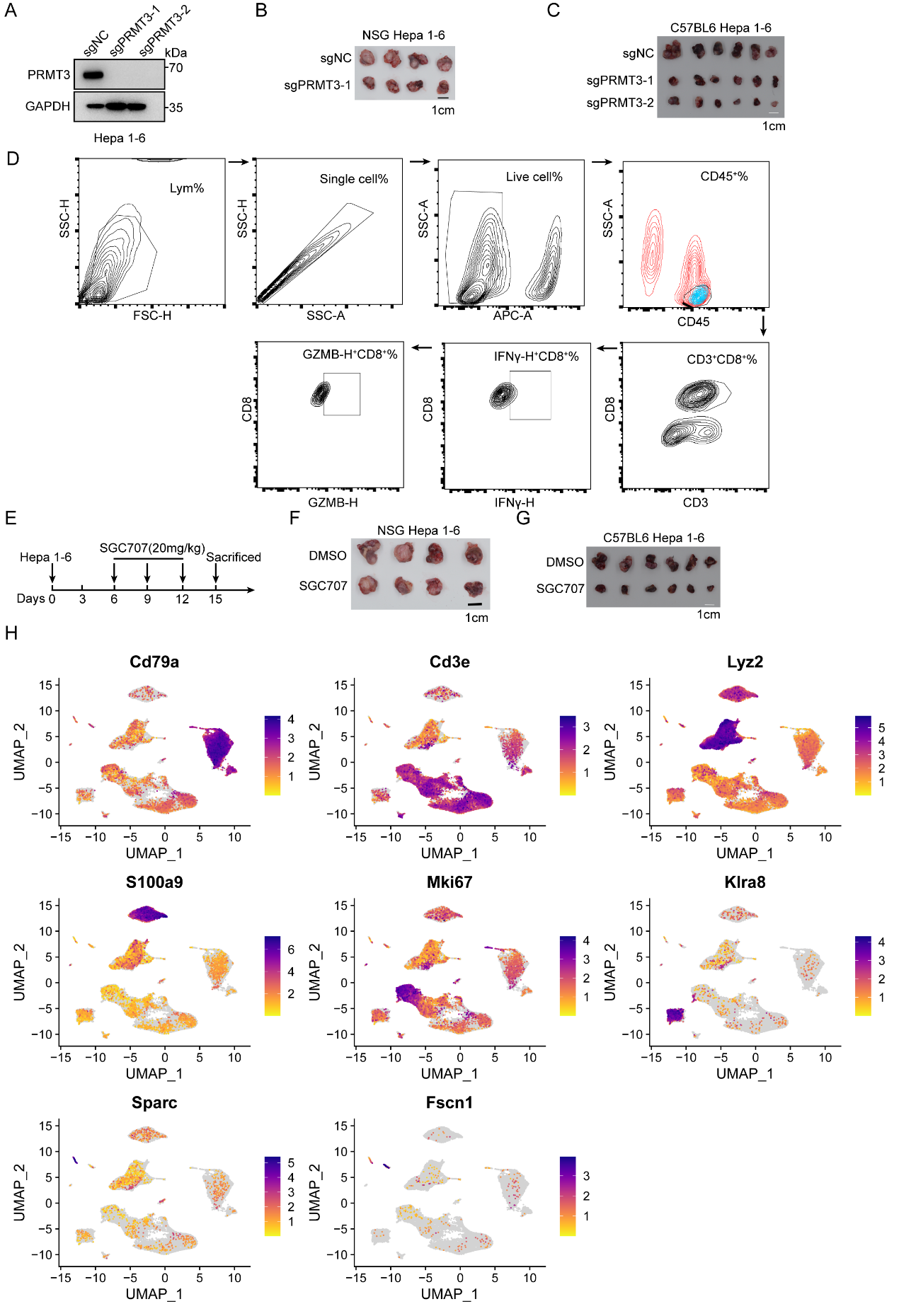

**Supplementary Figure** **5.** ***Prmt3*-KO or PRMT3 inhibition increases immune infiltration and activates T cell-mediated anti-tumor immunity**

A. PRMT3 expression was detected by western blot in *Prmt3*-KO and ctrl Hepa1-6 cells.

B. The effect of *Prmt3*-KO on subcutaneous tumor growth in NSG mice (n=5).

C. The effect of *Prmt3*-KO on subcutaneous tumor growth in C57BL6 mice (n=6).

D. Gating strategies for flow cytometry. Identification of CD8^+^T, IFNγ^+^ CD8^+^T cells and GZMB^+^ CD8^+^T cells.

E. Schematic diagram illustrates the workflow of Hepa1-6 subcutaneous model treated with DMSO or SGC707 (20 mg/kg) (n=6).

F. The effect of DMSO or SGC707 (20 mg/kg) on subcutaneous tumor growth in NSG mice (n=5).

G. The effect of DMSO or SGC707 (20 mg/kg) on subcutaneous tumor growth in C57BL6 mice (n=6).

H. Feature plot for clustering analysis in scRNA-seq.

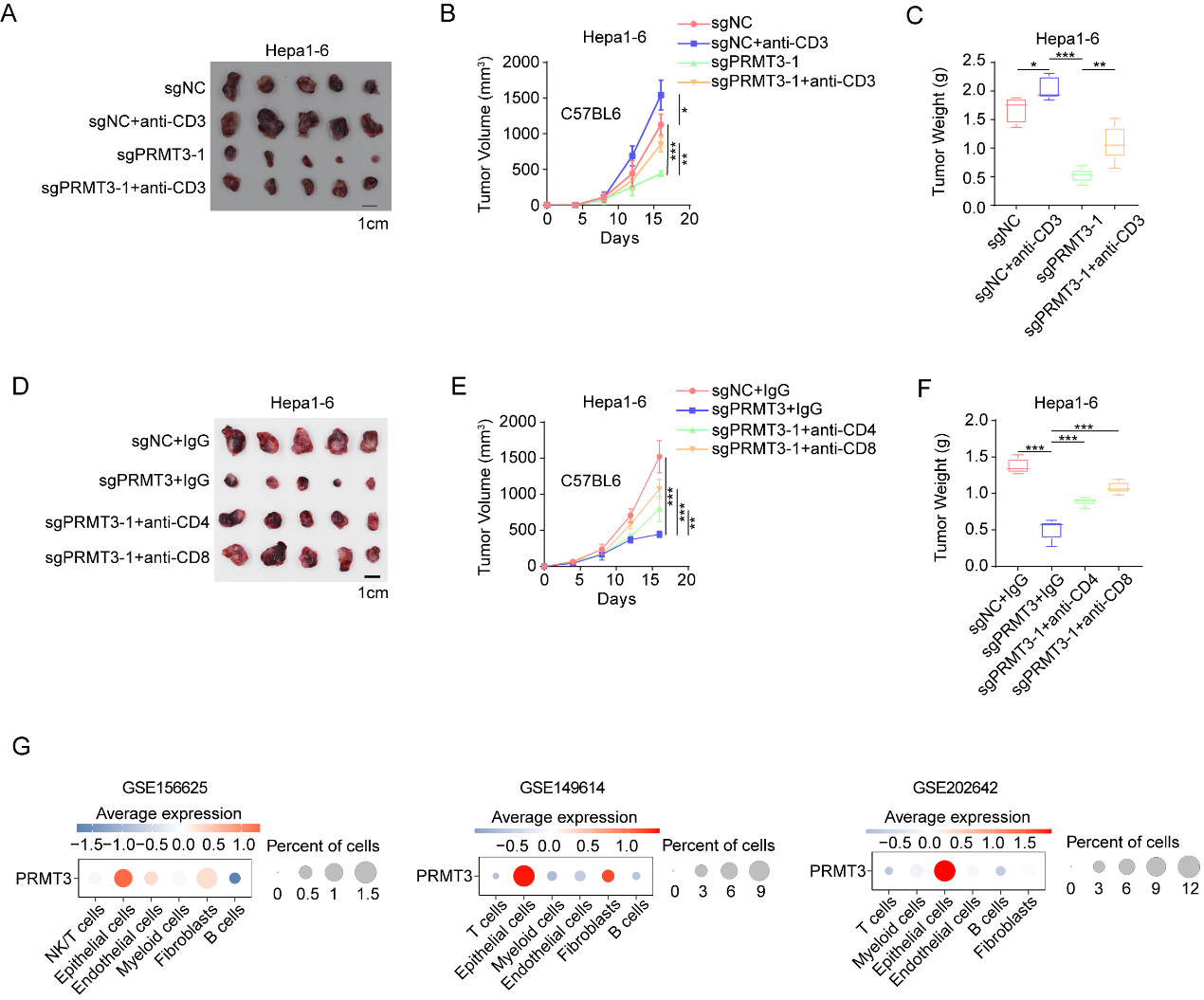

**Supplementary Figure** **6. T cells mediated the effect of PRMT3 on anti-tumor immunity in HCC**

A. The effect of *Prmt3*-KO and anti-CD3 antibody treatment on subcutaneous tumor growth in C57BL6 mice (n=6).

B, C. The measurement of tumor volumes (B) and tumor weights (C) of subcutaneously implanted Hepa1-6 cells (*Prmt3*-KO and *Prmt3*-WT) treated with anti-CD3 antibody in C57BL6 mice (n=5).

D. The effect of *Prmt3*-KO and anti-CD4/CD8 antibody treatment on subcutaneous tumor growth in C57BL6 mice (n=6).

E, F. The measurement of tumor volumes (B) and tumor weights (C) of subcutaneously implanted Hepa1-6 cells (*Prmt3*-KO and *Prmt3*-WT) treated with anti-CD4/CD8 antibody in C57BL6 mice (n=5).

G. PRMT3 expression in various cell types in the tumor niche was analyzed in several public dataset.

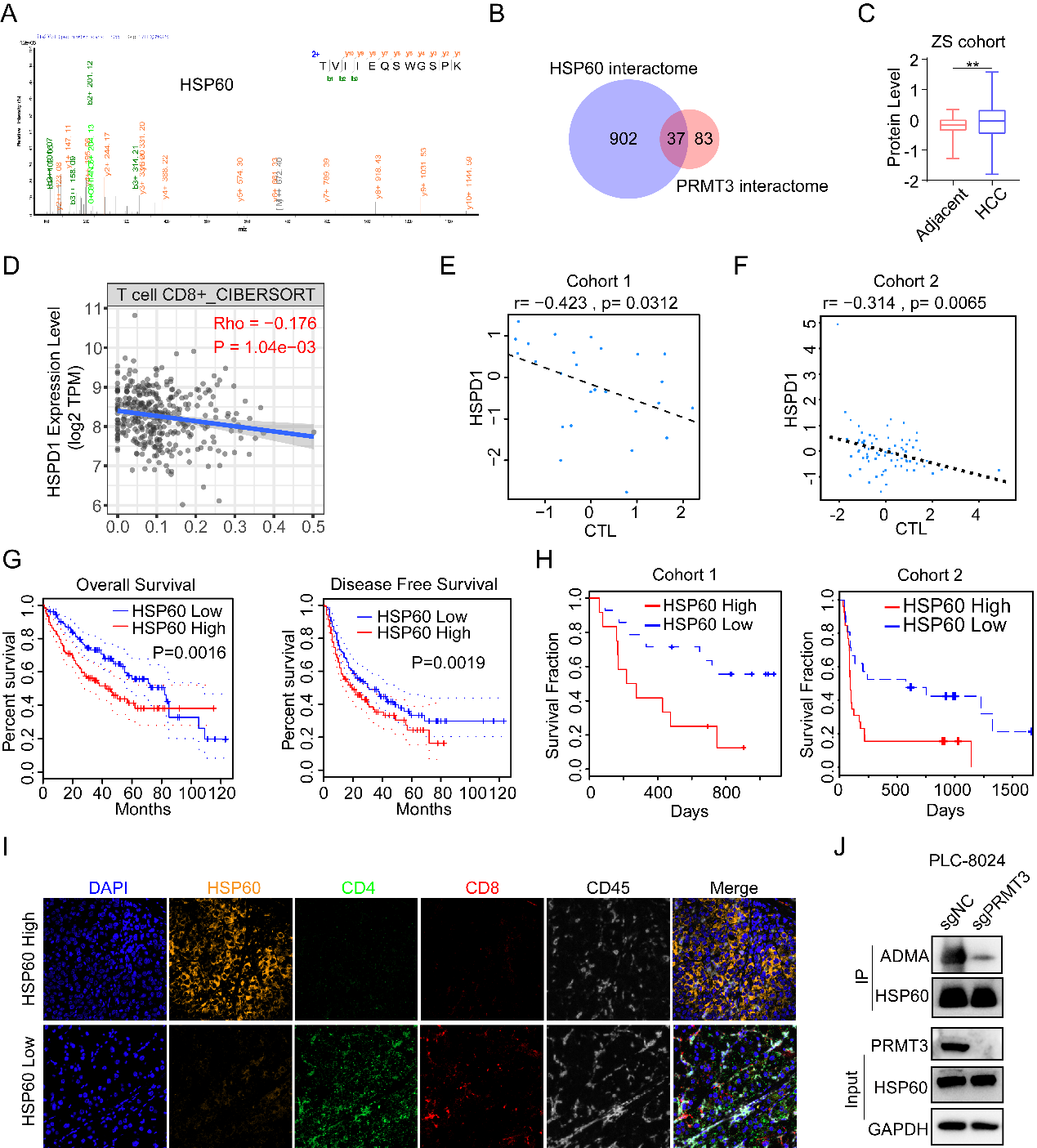

**Supplementary Figure** **7. PRMT3 methylates HSP60 at R446 and promotes its** **oligomerization**

A. Fragmentation spectrum of the HSP60 peptide identified by liquid chromatography/tandem mass spectrometry (LC-MS/MS).

B. Venn Diagram showing the 37(1/3) proteins were identified for both PRMT3 and HSP60 interacting proteins retrieved from the BioRID database.

C. HSP60 was significantly up-regulated in HCC compared to adjacent tissue in a public proteomics dataset.

D. Associations of HSPD1 expression with CD8^+^ T cell infiltration evaluated by CIBERSORT in TCGA-LIHC dataset.

E, F. Associations of *HSPD1* expression with CTL infiltration in two immunotherapy cohorts.

G. Kaplan–Meier overall survival curves of individuals with different *HSPD1* expression in TCGA-LIHC dataset.

H. Kaplan–Meier overall survival curves of individuals with different *HSPD1* expression in two immunotherapy cohorts.

I. Representative example of HSP60-high and HSP60-low HCC. Tumor staining by multiplexed IHC shows the spatial distributions of CD4^+^ T cells and CD8^+^ T cells.

J. WB analysis of immunoprecipitated HSP60 to determine the effect of *Prmt3*-KO on arginine methylation of HSP60 in Hepa1-6 cells using asymmetric dimethylarginine antibody (ADMA).

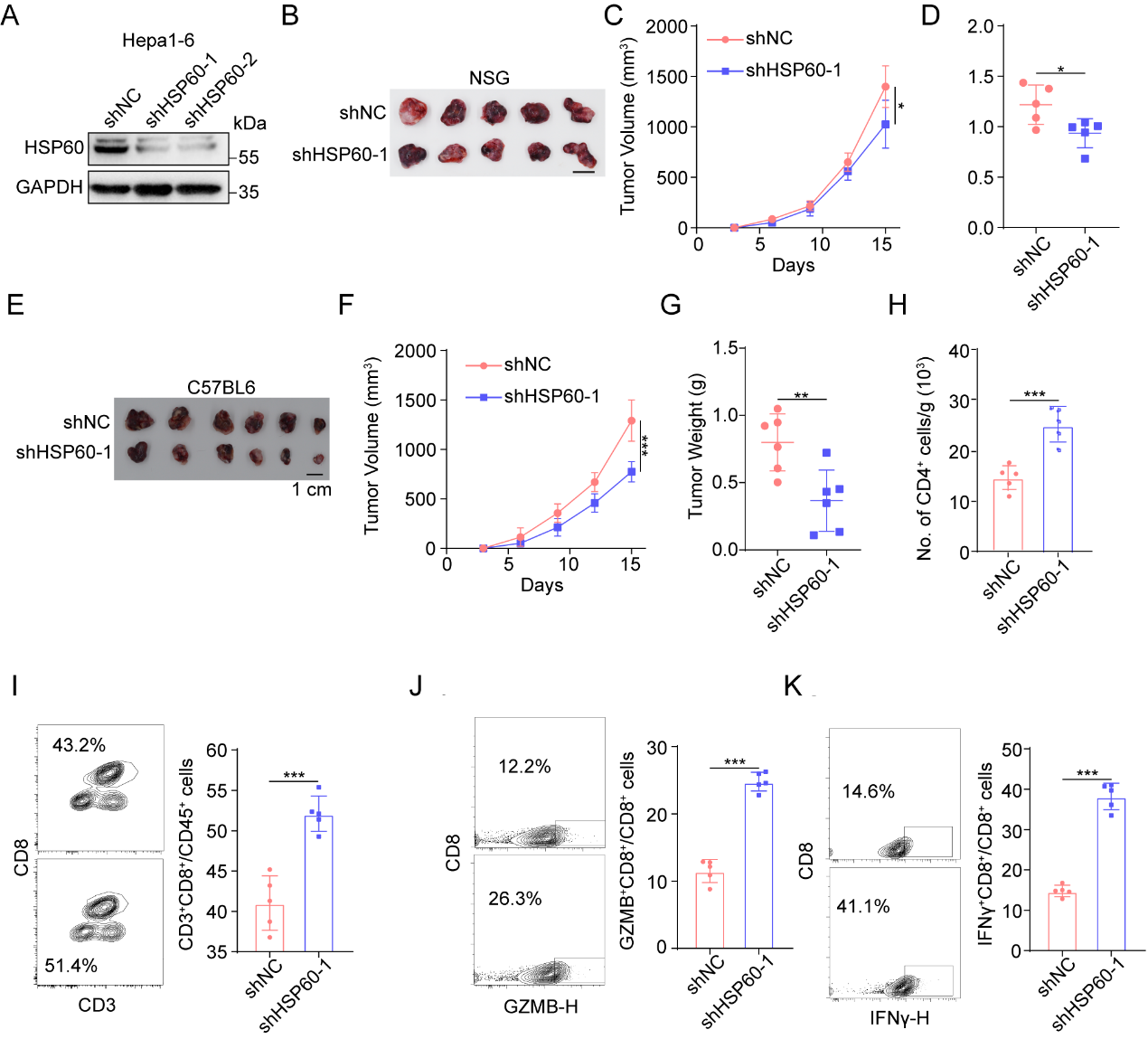

**Supplementary Figure** **8. HSP60 KD inhibited tumor growth and activated anti-tumor immunity**

A. HSP60 expression was detected by western blot in HSP60-KD and ctrl Hepa1-6 cells.

B. The effect of HSP60-KD on subcutaneous tumor growth in NSG mice (n=6).

C, D. The measurement of tumor volumes (C) and tumor weights (D) of subcutaneously implanted indicated Hepa1-6 cells in NSG mice (n=6).

E. The effect of HSP60-KD on subcutaneous tumor growth in C57BL6 mice (n=6).

F, G. The measurement of tumor volumes (F) and tumor weights (G) of subcutaneously implanted indicated Hepa1-6 cells in C57BL6 mice (n=6).

H, I. CD4^+^ (H) and CD8^+^ (I) T cells from tumors in ctrl and HSP60-KD groups were analyzed by flow cytometry.

J, K. Flow cytometry analysis assessing the percentage of T cell functional markers IFNγ (J) and GZMB (K) from tumors in ctrl and HSP60-KD groups.

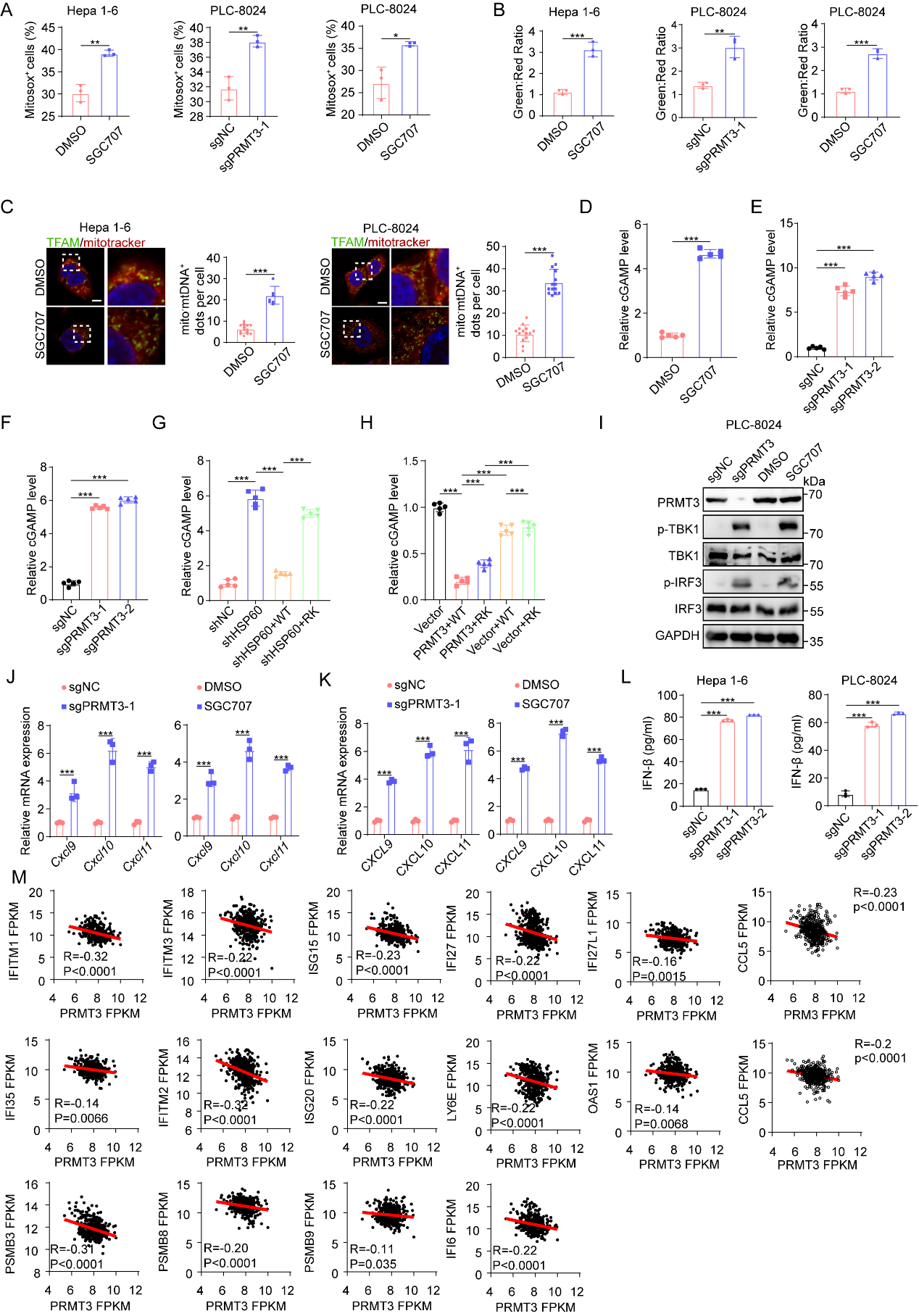

**Supplementary Figure 9.** **Inhibition of PRMT3-mediated arginine methylation of HSP60 induces** **mtDNA leakage and cGAS/STING signaling activation**

A. Mitosox+ cells were detected by flow cytometry in indicated groups.

B. JC-1 aggregates: JC-1 monomers ratio in indicated groups.

C. The effect of SGC707 treatment on mtDNA release was assessed by IF staining with mitotracker and TFAM. Representative images (scale bar, 5 μm) and quantitative results are shown.

D-H. cGAMP level was analyzed in indicated groups with ELISA assay.

I. The effects of *PRMT3*-KO or SGC707 treatment on expression of TBK1/p-TBK1 and IRF3/p-IRF3 in PLC-8024 cells.

J. The effects of *PRMT3*-KO and SGC707 treatment on expression of interferon-stimulated genes in Hepa1-6 cells.

K. The effects of *PRMT3*-KO and SGC707 treatment on expression of interferon-stimulated genes in PLC-8024 cells.

L. The effects of *PRMT3*-KO on IFNβ production in Hepa1-6 and PLC-8024 cells.

M. Correlation between several interferon-stimulated genes and *PRMT3* in TCGA-LIHC dataset.

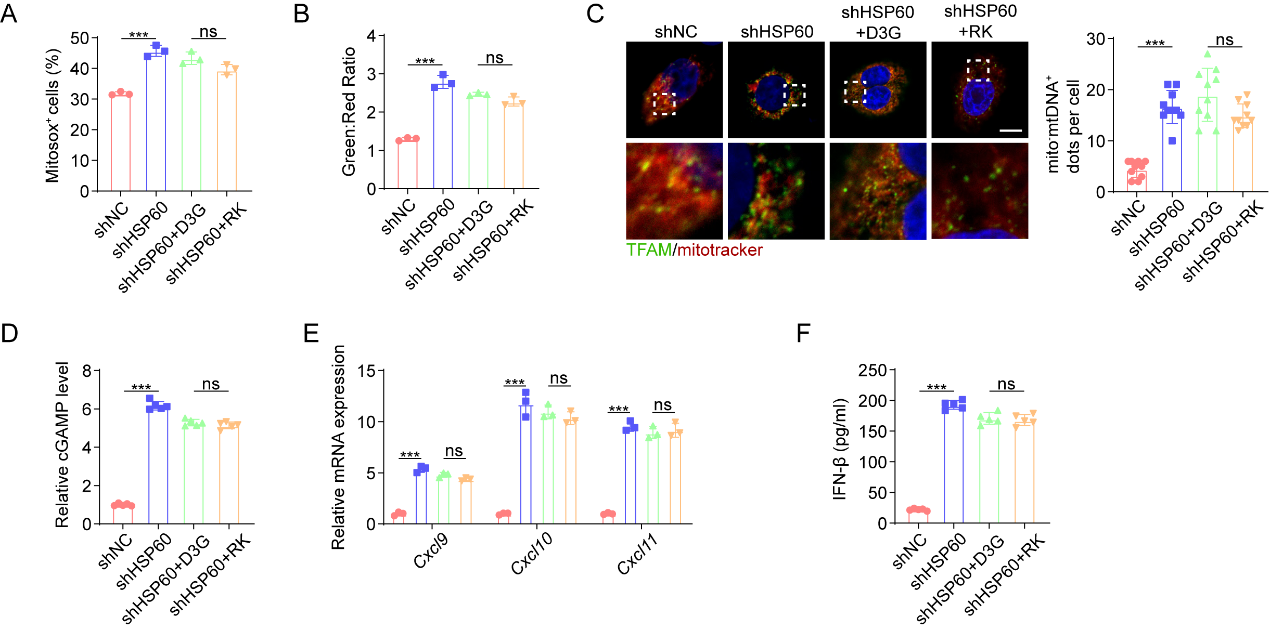

**Supplementary Figure 10.** **Both HSP60-D3G and HSP60-RK mutant induces mtDNA leakage and cGAS/STING signaling activation**

A. Mitosox+ cells were detected by flow cytometry in indicated groups.

B. JC-1 aggregates: JC-1 monomers ratio in indicated groups.

C. The effect of SGC707 treatment on mtDNA release was assessed by IF staining with mitotracker and TFAM. Representative images (scale bar, 5 μm) and quantitative results are shown.

D. cGAMP level was analyzed in indicated groups with ELISA assay.

E. The effects of HSP60-D3G and HSP60-RK mutant on expression of interferon-stimulated genes in Hepa1-6 cells.

F. The effects of HSP60-D3G and HSP60-RK mutant on IFNβ production in Hepa1-6.

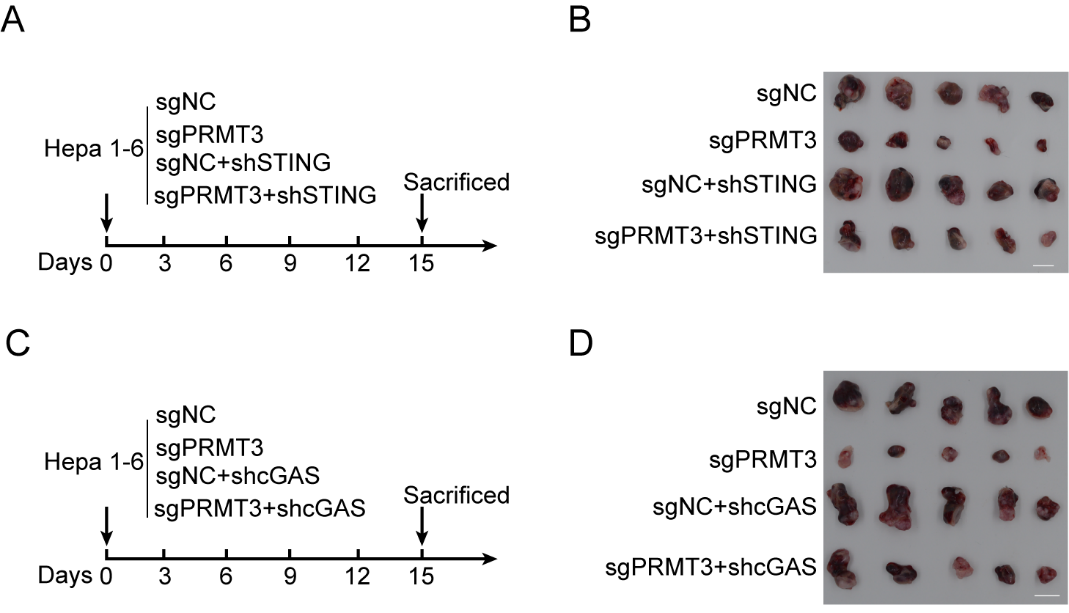

**Supplementary Figure 11.** **The anti-tumor immunity induced by *Prmt3*-KO or inhibition was dependent on the activation of cGAS/STING signaling**

A. Schematic diagram illustrates the workflow of subcutaneous model implanted indicated Hepa1-6 cells (n=5).

B. The effect of STING-KD on the tumor growth of subcutaneously implanted *Prmt3*-KO Hepa1-6 cells (n=5). Scale bars, 1 cm.

C. Schematic diagram illustrates the workflow of subcutaneous model implanted indicated Hepa1-6 cells (n=5).

D. The effect of cGAS-KD on the tumor growth of subcutaneously implanted *Prmt3*-KO Hepa1-6 cells (n=5). Scale bars, 1 cm.

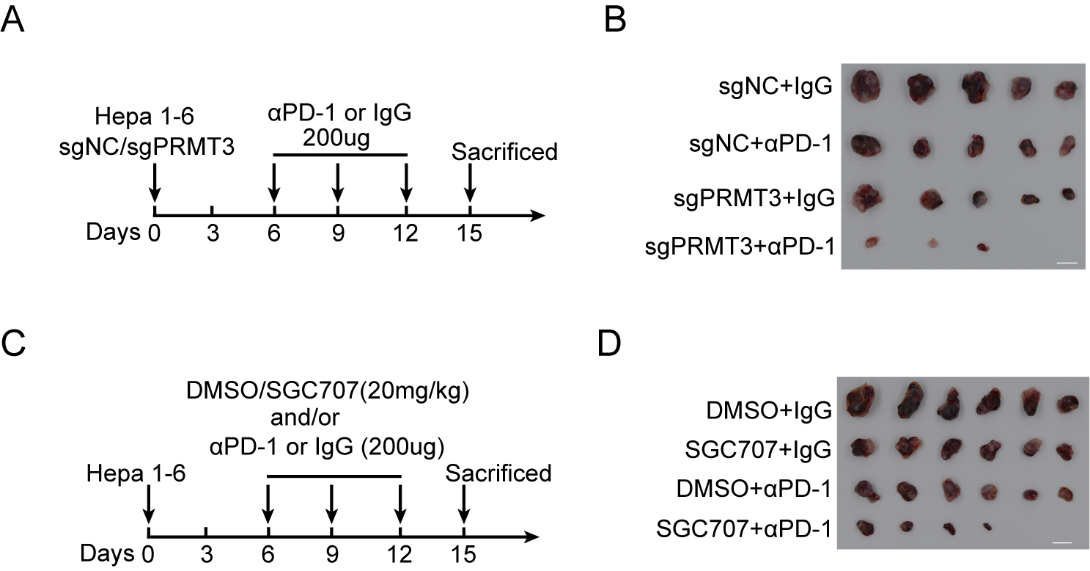

**Supplementary Figure 12.** ***Prmt3*-KO or inhibition enhances immunotherapy response in HCC**

A. Schematic diagram illustrates the workflow of subcutaneous model implanted indicated Hepa1-6 cells and treated with PD-1 antibody or IgG (n=5).

B. The effect of *Prmt3*-KO on the tumor growth of subcutaneous tumor growth treated with PD-1 antibody or IgG (n=5). Scale bars, 1 cm.

C. Schematic diagram illustrates the workflow of the subcutaneous model treated with SGC707 and/or PD-1 antibody (n=6).

D. The effect of SGC707 treatment on the tumor growth of subcutaneous tumors treated with PD-1 antibody or IgG (n=5). Scale bars, 1 cm.

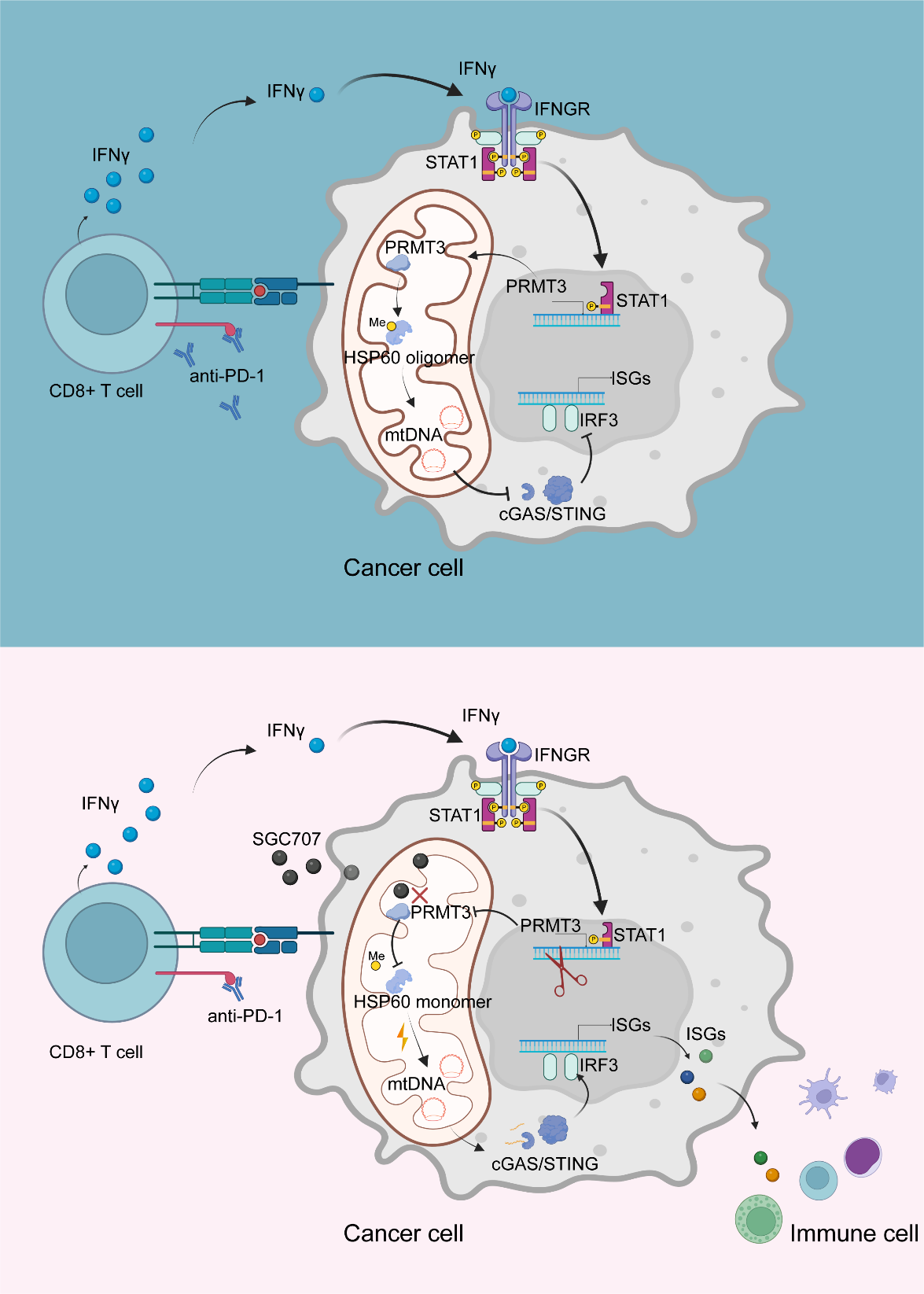

**Supplementary Figure 13.** Working model. Immunotherapy induces activation of effector T cells to trigger PRMT3 expression in cancer cells via an IFNγ-STAT1-dependent pathway. PRMT3 methylates HSP60 at R446 to induce its oligomerization and the maintenance of mitochondrial homeostasis. Inhibition of HSP60-R446 methylation disrupts mitochondrial homeostasis, which activates the cGAS/STING pathway and anti-tumor immunity.
